# Artificial selection for predatory behavior results in dietary niche differentiation in an omnivorous mammal

**DOI:** 10.1101/2021.11.15.468574

**Authors:** Anni Hämäläinen, Mikko Kiljunen, Esa Koskela, Pawel Koteja, Tapio Mappes, Milla Rajala, Katariina Tiainen

**Affiliations:** Institute of Environmental Sciences, Jagiellonian University, Cracow, Poland; Department of Biological and Environmental Science, University of Jyväskylä, Finland

**Author notes:** Corresponding author: ORCID: 0000-0001-9260-8299. Author contributions:* AH designed and performed the statistical analyses and wrote the paper. EK, PK and TM planned and designed the study. PK developed and provided the selection lines. EK and TM carried out the field study. MK designed and supervised the isotope analyses. AH, MR, and KT carried out the isotope analyses and food source sampling. All authors provided input for and approved the final manuscript.

**Keywords:** artificial selection, bank vole, diet choice, dietary niche, omnivory, predatory behavior, specialization, stable isotopes, trophic niche

## Abstract

The diet of an individual is a result of the availability of dietary items and the individual’s foraging skills and preferences. Behavioral differences may thus influence diet variation, but the evolvability of diet choice through behavioral evolution has not been studied. We used experimental evolution combined with a field enclosure experiment to test whether behavioral selection leads to dietary divergence. We analysed the individual dietary niche via stable isotope ratios of nitrogen (δ^15^N) and carbon (δ^13^C) in the hair of an omnivorous mammal, bank vole, from 4 lines selected for predatory behavior and 4 unselected control lines. Predatory voles had higher hair δ^15^N values than control voles, supporting our hypothesis that predatory voles would consume a higher trophic level diet (more animal vs. plant foods). This difference was significant in the early but not the late summer season. The δ^13^C values also indicated a seasonal change in the consumed plant matter and a difference in food sources among selection lines in the early summer. These results imply that environmental factors interact with evolved behavioral tendencies to determine dietary niche heterogeneity. Behavioral selection thus has potential to contribute to the evolution of diet choice and ultimately the species’ ecological niche breadth.

## Introduction

The diet an individual consumes is a result of the availability of different food items, the species-specific dietary range, and individual specialization [1,2]. The resulting differences among individuals in dietary niche are generated by environmental and genetic variation as well as phenotypic plasticity [1,2], but the relative effects of these factors have rarely been tested [2–5]. Inherited effects might manifest through various morphological, physiological, or behavioral traits that shape preferences for certain foods and specialized behaviors connected to seeking or processing food items [3]. For example, stable individual differences have been observed in hunting behaviors (antlions *Myrmeleon hyalinus* [6], guillemots *Uria lomvia* [7]). Genetically determined behavioral differences could thus contribute to dietary specialization at the individual level, with significant consequences for resource competition and even community functioning [2,8,9]. Yet, the role of evolved behavioral traits in shaping diet choice remains poorly understood [3] and research has focused primarily on predatory behavior of carnivores, which tend to show more specialization than other trophic groups [2]. Deciphering the relative effects of genetic and environmental influences on realized diets of individuals is challenging but essential for understanding the evolvability and plasticity of the dietary niche [5]. In this study, we address this problem by combining artificial selection for a predatory behavior in an omnivorous mammal with a field experiment.

Omnivores, animals that consume diets from more than one trophic level [10], are an understudied but exceptionally interesting group for diet choice studies because of the broad range of different types of dietary items they can potentially consume. This potential could facilitate trophic niche heterogeneity among individuals under intraspecific competition. The wide potential dietary breadth of omnivores involves morphological and physiological adaptations such as changes in dentition, gut length and structure, digestive enzymes and stomach acidity compared to related herbivorous or carnivorous species [10–13]. Individual-level variation in trophic niche has not been previously linked with behaviors, although such divergence could be widespread and under selection in omnivores. Behavioral adaptations such as capturing and processing prey would be required to transition from strict herbivory to omnivory. Yet, the significance and evolvability of behavioral traits associated with the diet breadth of omnivores remains unknown.

In this study, we assessed the significance of a behavioral adaptation on diet choice in an omnivorous rodent, the bank vole (*Myodes [Clethrionomys] glareolus*). We quantified the relative importance of a genetically determined behavioral type and environmental variation on the realized diet of individuals. We compared the dietary niche of bank voles from lines artificially selected for an increased predatory tendency [14] with unselected lines by measuring stable isotope ratios of carbon and nitrogen in the hair of field-reared individuals of both types. Stable isotope methods [15] are suited to studying individual dietary niches as they permit an evaluation of the consumed diet based on the isotopic signatures in the animal’s tissues, integrating dietary information over longer time periods. A higher isotope ratio of nitrogen (^15^N/^14^N) indicates consumption of food items from a higher trophic level because δ^15^N is enriched along the food chain [16]. Isotope ratios of carbon (^13^C/^12^C) in turn are more conserved through the food webs but variable among primary producers [16] and allow differentiating among consumers’ diet sources. Using these isotope ratios as indicators of long-term diet choice, we evaluated the potential for behavioral selection to shape the dietary niche. We specifically hypothesized that the artificially selected tendency for predatory behavior would lead to the consumption of a diet from a higher trophic level (higher proportion of animal sources, such as invertebrate prey) in field conditions, indicated by a higher δ^15^N ratio in the hair of predatory relative to control line voles.

## Materials and methods

### Study system

The bank vole is a common, widespread rodent, whose dietary profile is uniquely placed among European rodent species, occupying an intermediate niche between herbivorous arvicoline species and granivorous-insectivorous murine species [17,18]. The majority of their diet consists of different plant sources (seeds, leaves, flowers, roots, bark) [17,19–21], but the proportion of animal matter (primarily invertebrates) in stomach contents can range from 0-23% [21–24] and the proportion of fungi from 0-10% [18,21–25] among populations and seasons. The majority of the animal food consists of insect larvae especially in the early season, but adult insects, worms or molluscs and vertebrate remains are infrequently consumed [18,21,25]. Possible heterogeneity in diet among individual bank voles remains poorly known because the relative proportions of different dietary items consumed by individuals over time has been difficult to assess with gut content analyses (but see [20]). The degree of dietary niche divergence among individuals is therefore unknown.

To test the importance of artificial selection (overall genotypic differences) in the realized diet, we used bank voles from a unique long-term selection experiment (for details see [14,26,27]). Briefly, several selection lines were established from a source population of 320 voles captured in Poland in years 2000-2001. To generate voles with a “predatory” phenotype, bank voles have been selected for their propensity to rapidly capture a live cricket under standardized conditions. Four parallel predatory (P) and four unselected control (C) lines are maintained. The continued selection has resulted in significant divergence in predatory efficiency, with predatory voles catching the cricket more than five times more often than control voles by the 24^th^ generation [27]. In the present study, we used descendants (offspring and grand-offspring) of the 25^th^ selected generation. The voles used in this experiment were never exposed to live prey prior to the experiment. The parental generation (“founders”) were born and reared in laboratory conditions at the University of Jyväskylä, Finland with ad lib water and standard rodent chow (Avelsfoder för råtta och mus R36; Lactamin, Stockholm, Sweden; 301 kcal /100 g; macronutrient content: 18.5 % protein; 4.0 % fat; 55.7 % carbohydrate) until release to field enclosures.

### Field experiment

To test whether the voles selected for a predatory tendency consumed a diet from a higher trophic level than control voles, we performed a field experiment. Founder voles were released into eleven 0.2 ha field enclosures near Konnevesi research station in Central Finland over two replicate experimental rounds in early (June-July) and late (August-September) summer in 2018 (total 22 enclosure replicates). The field enclosures had early succession vegetation consisting primarily of grasses, forbs and shrubs. This study was performed in connection with a larger field experiment designed to test density- and frequency-dependent selection on behavioral tactics (TM et al. in prep.), for which the initial density (8 or 16 adults per enclosure) and ratio of the P- and C-line adults (1:3 or 3:1) varied among the enclosures. The initial adult sex ratio was 1:1 in all enclosures.

The founders were mated (maintaining selection line separation) in spring-summer 2018 in the animal facilities in Jyväskylä. Females were monitored daily to determine the exact date of delivery. Within a day of the birth of a litter, the pups were individually marked by distal phalanx removal, sexed, weighed, and their head widths measured. After parturition, the females with their newborn litters were transported into the enclosures in their home cage [28]. The cages were placed open and on their side on the ground with partial shelter and a small quantity of food (approx. two days minimum energy requirement) so that the females could transport the pups out at leisure. Litter size ranged 2-7 (C mean = 4.13, P mean = 4.47), with the total initial number of pups released per enclosure ranging 15-38.

The dams were left to rear the young to independence on a natural diet. After ca. 25 days, when the juveniles move around independently, all animals were captured from the enclosures using live traps baited with sunflower seeds and potato (details on enclosures and live trapping e.g. [29]). In total, 133 weaned young (65 P-line, 68 C-line) were captured from the enclosures in the two rounds (first: N=50, second: N=83). The number of weaned offspring per enclosure per round ranged from 0 to 28 individuals. One enclosure had no surviving offspring in either round and another two enclosures had none in the first round. Captured young voles were identified, sexed, weighed, head width measured, and a small patch of hair was clipped from the back with scissors (aiming to collect entire hair shafts) for isotope analyses. All hair samples for isotope analyses were thus derived from individuals that had spent their entire lifetime, from age 1-3 days (i.e. before growing any fur) until sampling, in the field enclosures. The dams relied on natural food items after the first few days of lactation. The pups begin to feed on solid food by the age of ca. 2 weeks (personal observations from lab conditions) and are weaned by age 20 days [14,30]. Thus, the isotope composition in the hair of the weaned juveniles consists of the combined effects of the diet consumed by the individual and by its mother. The hair samples were stored in Eppendorf tubes in room temperature until analyses in 2019 summer.

### Collection of possible dietary items

To relate the isotope ratios in the voles’ hair to the available food items, we collected samples of plants, invertebrates and fungi from the field enclosures and analysed their isotope signatures. The detailed methods are provided in the Supplementary Information (SI).

### Isotopes of captive voles

To account for the possibility that any differences between the selection lines are due to intrinsic differences in physiology (e.g. differential fractionation into hair due to differences in metabolism), we collected hair samples from individuals that had lived their entire lives in the lab on the standard rodent diet supplied to the adults in this experiment before their release into the field enclosures. We shaved hair from the backs of two females from each of the four parallel predatory selection lines and the four control lines, producing eight samples per selection direction (total N=16). The samples were analysed in the same manner as the samples derived from the field conditions.

### Isotope analyses

Lipids were removed from the hair samples with a Chloroform-Methanol extraction [31], samples were dried and then 0.5-0.7 mg of each sample was weighed into tin capsules for isotope measurements. All samples representing vole diet (invertebrates, plant material, fungi) were freeze-dried to a constant weight, ground to a fine powder using a ball mill or mortar and pestle, and then also weighed into tin capsules. Stable isotope analyses for carbon and nitrogen were conducted using a Thermo Finnigan DELTA_plus_ Advantage continuous-flow stable isotope-ratio mass spectrometer (CF-SIRMS) coupled with a FlashEA 1112 elemental analyzer. Results are expressed using the standard δ notation as parts per thousand (‰) differences from the international standard. The reference materials used were internal standards of known relation to the international standards of Vienna PeeDee Belemnite (for carbon) and atmospheric N_2_ (for nitrogen). Precision was always better than 0.13‰ for carbon and 0.38 for nitrogen, based on the standard deviation of replicates of the internal standards.

### Fractionation coefficients

To relate the stable isotopes in vole hair to the isotope ratios of possible food items, we determined the average fractionation, i.e. difference in isotope ratios between the consumed food items and the measured isotope ratios in hair. We used the 16 samples collected from captive voles maintained on a standard diet of rodent pellets to determine the degree of fractionation. We computed the average isotope values for the rodent pellets fed to the captive voles as δ^15^N = 1.778±0.268 (mean±SD), and δ^13^C = −26.613±2.491. We related these to the isotope values measured from the hair of the captive voles and determined the trophic enrichment factors (TEFs, Δ) as Δ^15^N = 5.335±0.553, and Δ^13^C = 2.145±0.239. These values were used to correct the isotope ratios of the food source samples from the field experiment to associate the food items with the vole isotopes.

### Statistics

Inspection of the isotope data indicated an outlier in δ^15^N (δ^15^N=8.73, 4 SD divergence from mean δ^15^N; Grubbs’s outlier test: G = 5.12996, U = 0.79912, P <0.001) that skewed the δ^15^N data disproportionately. As the reason for the exceptionally high reading was unknown but might indicate e.g. a sample handling error, we chose to conduct all further analyses without this observation, with a final sample size of N=132 for all analyses. Including the outlier in the models did not qualitatively change the analysis outcomes but reduced the significance of some results (SI, Table S2). The isotope data with both experimental rounds combined were not normally distributed (Shapiro-Wilk test of δ^13^C and δ^15^N both P<0.001), so we used Wilcoxon tests for bivariate analyses of the raw data. Differences in dietary variation (i.e. individual niche differentiation) between seasons and selection lines were explored with Levene’s Test for Homogeneity of Variance (median-centered approach; *car*-package [32]).

We constructed linear mixed effects models (LME) to examine the effects of selection and environment on isotope values while accounting for maternal effects and pseudoreplication. Diagnostic plots of the initial models with untransformed data indicated possible problems with the normality assumption. We therefore performed Box-Cox power transformations to the isotope data distributions (packages forecast [33,34], EnvStats [35]) and selected the best approximations for normality. For δ^15^N, transformation was done with λ=-1, for δ^13^C we first transformed all values to positive ((−1*min(δ^13^C))+1) and then applied Box Cox transformation with λ =2. We then used a Gaussian error distribution with an identity link function for both models. Fulfilment of model assumptions was assessed by visual inspection of qq-plots, observed values vs. residuals and leverage of specific data points. While qq-plots suggested the presence of some outliers in the residuals, these data points did not have a high leverage on the estimated model, thus all data were retained in the models. The results for both δ^13^C and δ^15^N were qualitatively robust when computing the same models using untransformed data (SI Table S2). We present the model-derived estimates for the Box-Cox-transformed data and back-transformed estimates for the variables of interest.

For each response variable (δ^15^N and δ^13^C values in hair), we created a full LME-model including as predictor variables the selection regime (C=control, P=predatory), experimental round (1=early, 2= late summer), density treatment (high, low), sex (male, female), and body condition (residual body mass relative to head width). We also included an interaction term of selection regime and round to test for the possibility that the seasonal food availability would influence the realized diets of the selection regimes differently. As intraspecific competition is thought to increase selection for niche divergence [2,36–38] we also evaluated the possibility that niche divergence between the lines is higher in high-density conditions by including an interaction term of density treatment and selection regime. When an interaction term was statistically non-significant (P>0.05), it was dropped from the model to facilitate easier interpretation of the main effects. In all models, we included the random effect of the enclosure (1-10; possible differences in microhabitat and in the social environment) and mother’s identity (N=52).

All analyses were completed using R program version 4.0.3 [39]. We fitted LMEs with restricted maximum likelihood estimation using the R-package lme4 [40]. P-values were computed using Satterthwaite’s method with the package lmerTest [41]. Marginal and conditional pseudo-R^2^-values were computed using MuMIn package [42]. The results were visualized using packages ggplot2 [43], ggsignif [44], sjPlot [45].

## Results

In the laboratory, isotope ratios of captive voles indicate no significant difference between the predatory and control voles in δ^13^C (W = 43, P= 0.279) or δ^15^N (W = 50, P = 0.065; Figure S1). Thus, any differences between the lines observed in the field conditions are likely not due to intrinsic differences in physiology (e.g. differential assimilation of macronutrients).

The δ^15^N values were strongly affected by an interaction between the effects of selection regime and the experiment round (Figure 1A, Figure 2; Table S1). In line with our hypothesis, the δ^15^N values of the predatory selection direction were higher than those of the control-line voles, indicating that voles selected for predatory behavior consumed a diet from a higher average trophic level than non-selected voles. The back-transformed estimates indicate a ca. 12 % difference in δ^15^N between C and P regimes in the early season (predicted δ^15^N for C: 5.88 [95% CI: 5.88, 6.25]; for P: 6.67 [6.25, 6.67]). However, in the late season no difference between the selection regimes was found (predicted δ^15^N for C: 5.88 [5.56, 6.25]; for P: 5.88 [5.56, 5.88]). δ^15^N values were on average slightly higher in the low-density treatment, suggesting that higher intraspecific competition may lead to an overall lower-level dietary niche. This effect of density was not dependent on selection regime (interaction of density and selection P>0.1).

**Figure 1.**
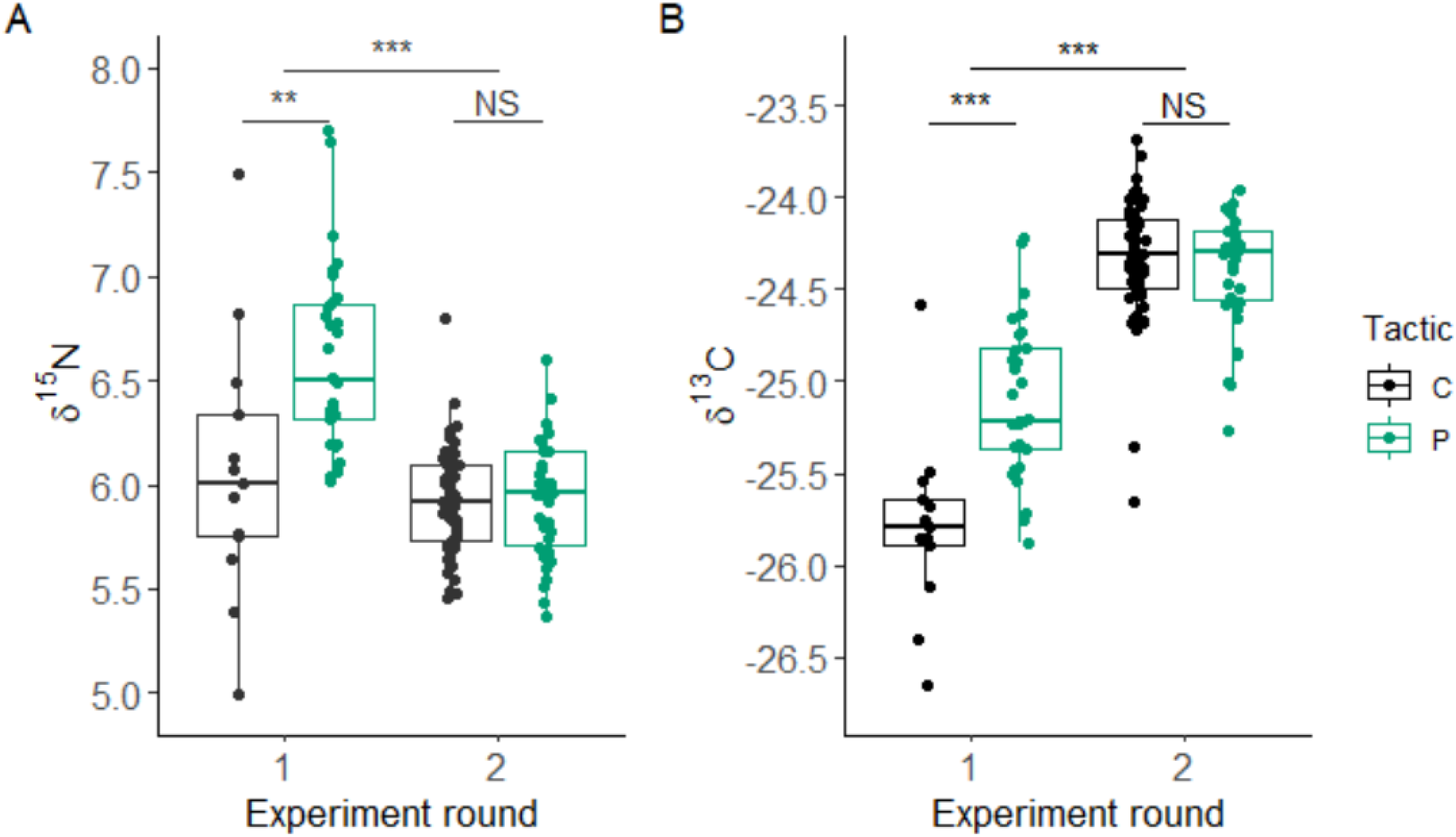
Raw data distribution for A) δ^15^N and B) δ^13^C for the two experimental rounds (1= early summer, 2= late summer) and the two selective regimes (P= predatory, C= control). N=132 after removal of one outlier (δ^15^N= 8.73 in experiment round 2, P-line vole). Asterisks indicate significant differences among groups based on Wilcoxon tests (NS: P>0.05, **: P=0.001-0.01, ***: P<0.001, details in SI).

**Figure 2.**
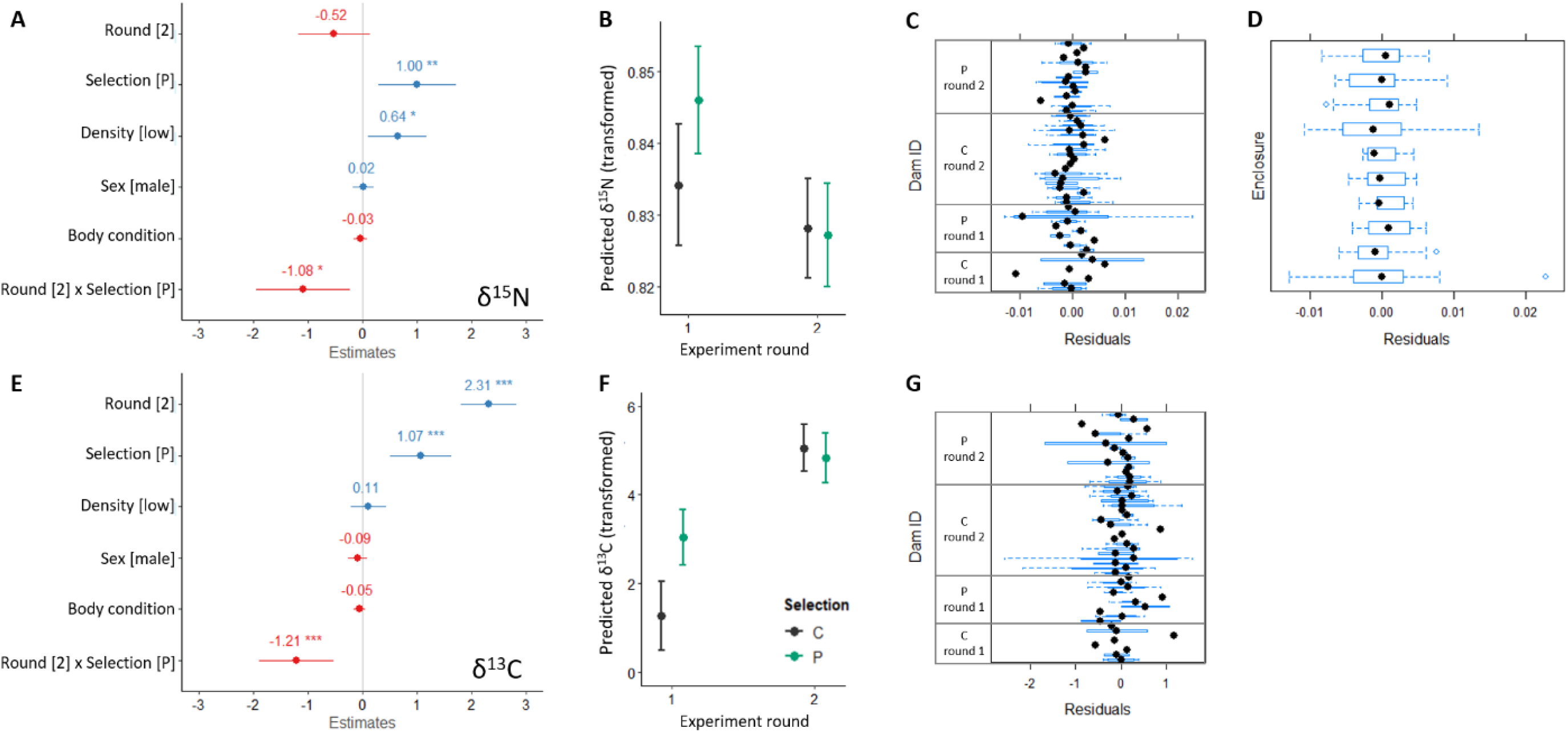
Effects of predictor variables on isotope ratios of δ^15^N (A-D) and δ^13^C (E-G). Forest plots for A) δ^15^N and E) δ^13^C show standardized (divided by 2 SD) estimates for fixed effects derived from linear mixed-effects models (SI Table S1) with Box-Cox-transformed isotope ratios as the response variables. The dots and associated numbers denote the relative effect, horizontal lines indicate 95% confidence intervals and asterisk notation refers to the effect significance (P-value). The predicted values and 95% CI for the marginal effects of the experimental round * selection regime interaction from these models are shown in panel B) for δ^15^N and F) δ^13^C. Intercepts and residual variance of the random effects are shown for C) mother’s id and D) enclosure for the δ^15^N model, and G) mother’s id for δ^13^C. The grouping of selection regime (C, P) and experimental round (1,2) are shown for dams in figures C and G for convenience. See SI for further details.

Similarly, the δ^13^C-levels were higher in the predatory lines in the early but not in the late summer (Figure 2; Table S1). The back-transformed estimates indicate a ca. 3% difference in δ^13^C between C and P regimes in the early season (C: −25.77 [95% CI: −26.24, −25.39]; for P: −24.99 [−25.23, −24.77]) and a 0.2 % difference in the late season (C: −24.32 [−24.48, −24.16]; for P: −24.39 [−24.56, −24.22]). The δ^13^C-values were also significantly higher in the second replicate overall, suggesting that the voles’ diet likely consisted of different plant sources in early and late season (Figure 1B; Table S1). Density treatment did not influence δ^13^C (interaction with selection regime and main effect of density both P>0.1).

The variances of isotope ratios for raw data were significantly higher in the early summer than in the late summer for both δ^15^N (F_1,130_= 22.479, P<0.001; Fig 1A) and δ^13^C (F_1,130_=15.548, P<0.001; Fig 1B), suggesting an overall higher degree of niche differentiation among individuals in the early relative to late summer (see also SI Fig. S2). When split by selection regime, the seasonal differences remained significant for both selection regimes for δ^15^N (Control: F_1,66_= 12.196, P<0.001; Predatory: F_1,62_= 4.461, P=0.039) but not for δ^13^C (Control: F= 0.879, P=0.352; Predatory: F= 3.245, P=0.076). There were no significant differences in variance between selection lines in δ^13^C (P>0.1 overall and when split by season). For δ^15^N, variance was significantly higher overall for Predatory relative to Control line voles (F_1,130_=6.137, P=0.015), but this difference did not hold within seasons. Variances of δ^13^C or δ^15^N did not significantly differ among density treatments in either P or C voles.

Vole isotope values were mainly within the isotope bi-plot area bounded by the TEF-corrected dietary source values (Fig.3). As the isotope ratios of the voles are derived from the combination of the different food items they consumed, these results imply that the voles consumed a primarily herbivorous diet (δ^15^N values low relative to animal sources), with the higher δ^13^C values especially in the late season suggesting a high consumption of grass inflorescences and seeds and possibly fungi. Although the vole hair samples fall within the range of isotope values of the sampled food items, the slight bias towards the lower right corner suggests a possibility that some possible food sources were missed from the analyses (e.g. lichens [18,21,23] with high δ^13^C and low δ^15^N [46] were not encountered during sampling). This hampered the use of stable isotope mixing models (e.g. MixSIAR) to formally estimate dietary proportions (analyses not shown).

**Figure 3.**
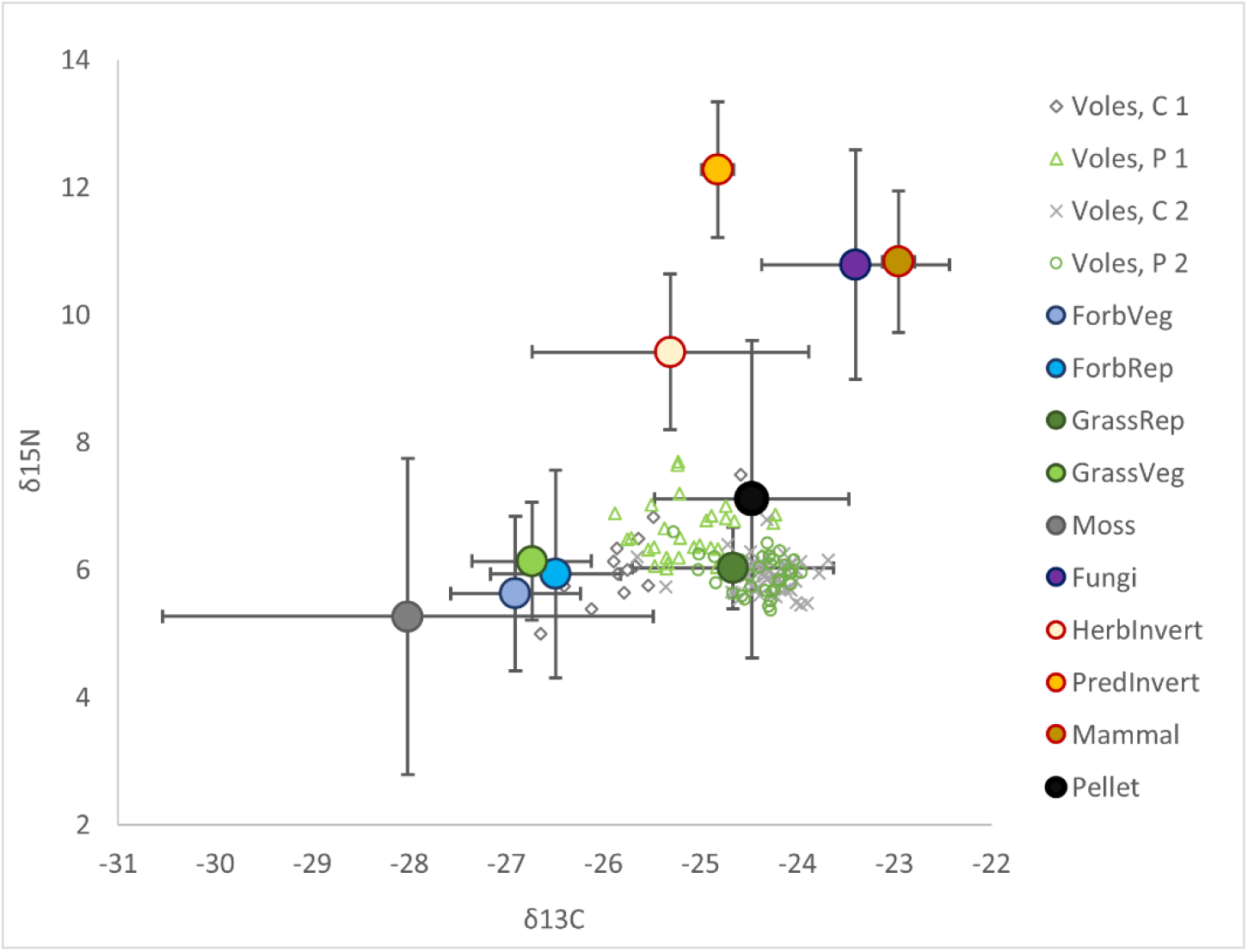
Isotope values of food sources and voles. Shown are mean values and standard deviations for vegetative parts (leaves, stems, roots) and reproductive parts (inflorescences, seeds) of forbs and grasses, mosses, fungi, herbivorous and predatory invertebrates, and mammal tissue (vole brain and muscle, shrew muscle). The isotope values of the food sources have been corrected for TEF (Δ^15^N: 5.335, Δ^13^C: 2.145). Vole hair values are raw data (experimental rounds 1 and 2, selection regimes C: control, P: predatory; see SI Figure S2 for the distribution of the vole hair samples only).

## Discussion

### Interaction of evolved behavioral type and environment create dietary niche variation

Individual dietary preferences are frequently observed, but the significance of a genetic component in foraging behaviors and thus the heritability of the dietary niche remains unresolved. Individual heterogeneity in dietary niche of omnivores and especially diet partitioning among different trophic levels is largely unknown despite the potentially significant implications for community functioning. This study provides the first evidence of an inherited foraging behavior affecting the niche divergence of an omnivorous rodent in a field setting. As predicted, an artificially selected predatory tendency was associated with a higher trophic level diet (higher δ^15^N isotope ratio suggesting consumption of more animal food relative to plants) relative to control line animals inhabiting the same field enclosures. This difference was limited to the early summer season, however, and disappeared later in the season, suggesting plasticity in dietary niche despite an underlying genetic tendency for dietary divergence. The diets of the predatory and control line animals also diverged in terms of carbon isotopes in the early summer, confirming that the realized diets of the selection lines differed. We conclude that differences in an inherited behavior were associated with a small but significant effect in dietary niche differentiation in interaction with the environment.

### Behavioral selection contributes to dietary niche differentiation

The genetic basis of individual niche variation remains poorly understood [2] but variation in realized diet may depend on e.g. preference, capacity to capture or process certain items, competitive ability, and behavioral types, which all have a genetic component (reviewed in [1–3]). Many foraging behaviors such as prey recognition or preference [4,47,48] as well as morphological traits relevant to prey selection are partly heritable [49], including the predatory tendency selected for in the vole selection lines used in this study [14,27]. This study shows that artificial selection for a behavior contributes to dietary niche divergence under natural food conditions. Interestingly, a complement of this association was found in sea otters [38], in which dietary specialization (due to food limitation) had consequences for behavioral phenotype divergence. Together, these studies indicate a possibly bidirectional association between diet and behavior. This first evidence of behavioral selection generating variation in diet suggests an intriguing prospect for a broader role of behavioral evolution contributing to dietary niche differentiation. Future studies should evaluate the possibility that selective pressures acting on behavioral syndromes [50] or traits such as risk taking, exploration and aggression could simultaneously impact on the niche breadth or specialization of individuals, contributing to associations between ecological roles and behavioral types.

Diet could be further shaped by other traits coevolving with the selected behavioral traits. Consistent behavioral traits frequently correlate with physiological or life-history traits that facilitate adaptation to specific environments [51–53]. For example, individuals with active personalities are expected to have a high metabolism and a high energy requirement, which in turn should be met by higher energy consumption, and possibly a broader dietary niche [3]. Thus, associations between dietary preferences and behavioral traits may be reinforced e.g. by the differing energetic needs and digestive efficiency associated with behavioral types [54–56]. Several other traits have been indirectly selected alongside the directional selection for an increased predatory tendency and prey catching speed in our study system [14,27]. Predatory lines tend to have a proactive behavioral style [26], possible stress sensitivity [27], and tendencies for aggression and an elevated sensitivity to hunger (according to transcriptome analysis [57]). The predatory phenotype is thus characterized by various mechanisms that can drive predatory foraging behaviors.

In addition to directly selected behaviors, niche divergence could also be facilitated by behavioral plasticity in diet choice and foraging [36,58]. Dietary flexibility itself can improve fitness [59] and if selection operates on genes that increase plasticity per se [60], behavioral plasticity in foraging could be under selection. Such plasticity may lead to individuals consuming a wider diet breadth or specializing in a broader range of different foods. Predatory voles in this study tended to have overall higher trophic niche heterogeneity, indicated by their higher variance in δ^15^N relative to control animals, and might exhibit higher dietary plasticity. Whether the plasticity results from higher specialization remains speculative because trophic niche position was determined from a single sample per individual, preventing assessment of within-individual diversity or consistency of diet choice. Specialization can have fitness benefits through the improved ability to effectively exploit certain resources [61–63] but entails possible trade-offs because of the limited flexibility in dietary range or foraging behaviors [3,64] (see also [59]). Specializing could also allow individuals to escape direct competition (e.g. switching to hunting instead of competing for plant protein), but we found no evidence of higher competition (density) influencing the dietary niche of predatory voles more than control voles.

The genetic component of the dietary niche development may be reinforced by cross-generational transmission of preferences in species with parental care. This possibility is suggested by the observed maternal effects in the isotope profiles of the juveniles (random effect of mother’s id), which might result from maternal genetic and epigenetic effects, a direct influence of the maternal diet choice through milk, or preferences or skills the young voles learned from their mothers. Juvenile nutrition during nursing is derived from maternal diet choices (guided by their genetic background) in the form of milk. The resulting isotope profile may be finetuned by the fractionation in isotopes between mother’s diet, isotope ratios in milk, and consolidation in offspring tissues. The foraging behaviors of the juvenile voles themselves may develop in part through observation and exposure to specific foods in early life (described for sea otters [65]). Notably, our sample captures the dietary niche variation of surviving young voles only and we do not have information on the diets of those voles that died early in the experiment. Dietary niche variation can have fitness effects (e.g. in pigeon guillemots [61], isopods [62], toads [59], and insect herbivores [66], see also [5]), thus the observed niche variation could result from the selective survival of those individuals that were able to best adapt their diet to the environment and intraspecific competition.

### Niche divergence is tempered by the environment

Features of the physical and social environment define what food resources are available to individuals. We observed overall seasonal differences in the isotope ratios in the hair of voles, likely due to the phenology of plants, animals and fungi altering the availability of specific dietary items over the summer. Energy-dense seeds are a preferred food [67] and seed abundance is lowest in the early summer, which might limit total energy availability. For example, in German farmland, animals and green plants made up the majority of bank vole diet in early summer, whereas cereals, seeds and fruits were consumed in later summer [20]. Stomach content analyses suggest that animal matter consumption typically peaks in the summer months [21–24]. Assuming a similar pattern at our study site, the early-season difference among the selection regimes could be explained for example by a dietary bias towards insect larvae by predatory lines.

The niche divergence among regimes and higher variance among individuals in the early season might thus be explained by stronger competition for preferred food items when seeds are unavailable. Individuals are thought to benefit from specializing to avoid competition when food is limited [1,63]. In this study, high-density treatment did not seem to increase specialization (variances did not differ between density treatments), but high density was associated with a lower overall trophic niche, possibly implying higher competition for food items from a higher trophic level in both P and C lines. Intraspecific competition for resources can lead to dietary niche expansion (e.g. broader niche use through behavioral changes in stickleback fish [58]) or to a narrowing niche (specialization [68]), depending on the environmental conditions [2] and genetic variation of the population [36]. Specialization has been observed on one hand when a broader range of food resources is available (e.g. fruit bats specializing more in specific fruits in a season when many different plants produce fruit [68]); on the other hand, generalist species may specialize more under poor nutritional conditions [69]. The specific outcome of the environment and intrinsic mechanisms of niche divergence can have significant effects on the stability of ecological networks [1,3].

### Conclusion

Given the possible genetic basis and potential fitness benefits of dietary niche flexibility or specialization under resource competition [36,59], traits associated with diet choice may be important targets for selection. We demonstrated that artificial selection for a predatory behavior shapes the diet of an omnivorous rodent in field conditions by increasing the predatory individuals’ trophic level. The dietary niche of individuals measured in the long term via stable isotope in hair indicated a small but consistent difference in dietary niche in interaction with the environment. Behavioral selection could, therefore, play a role in defining the trophic niche of individuals. Individual differences in diet choice and diet breadth can, in turn, have significant ecological consequences [1,3]. Our results point to the necessity of considering the significance of consistent behavioral variation in foraging when assessing the overall role of omnivores in the ecological community.

## Ethical statement

The research was conducted in accordance with the relevant laws and all procedures performed on the animals had an ethical committee approval (ESAVI/3981/2018).

## Open data statement

Raw data will be deposited in dryad upon acceptance for publication.

## Acknowledgements

We thank Tanja Hirvonen, Tommi Vuori and Konnevesi Research Station staff for help with field and laboratory work.

