## Supplementary information 2 for "Artificial selection for predatory behavior results in dietary niche differentiation in an omnivorous mammal"

| Kingdom | Group | Taxon | Type | Parts | Year | Round | d13C | d15N | %C | %N | C/N |
|---|---|---|---|---|---|---|---|---|---|---|---|
| Animal | Invertebrate | *Auchenorrhyncha* | Herbivorous invert. | whole body | 2019 | 1 | -27.30 | 1.74 | 53.40 | 9.83 | 5.43 |
| Animal | Invertebrate | *Auchenorrhyncha* | Herbivorous invert. | whole body | 2019 | 2 | -27.84 | 2.41 | 52.05 | 10.13 | 5.14 |
| Animal | Invertebrate | *Chrysomelidae* | Herbivorous invert. | whole body | 2019 | 1 | -27.60 | 4.36 | 49.37 | 10.03 | 4.92 |
| Animal | Invertebrate | *Chrysomelidae* | Herbivorous invert. | whole body | 2019 | 2 | -27.04 | 5.56 | 50.60 | 8.76 | 5.78 |
| Animal | Invertebrate | *Coleoptera* larvae | Herbivorous invert. | whole body | 2019 | 1 | -27.95 | 4.01 | 50.21 | 8.28 | 6.07 |
| Animal | Invertebrate | *Gastropoda* | Herbivorous invert. | whole body | 2019 | 2 | -28.52 | 3.21 | 43.50 | 2.12 | 20.47 |
| Animal | Invertebrate | *Gastropoda* | Herbivorous invert. | whole body | 2019 | 2 | -23.16 | 4.56 | 29.07 | 4.56 | 6.37 |
| Animal | Invertebrate | *Heteroptera* | Herbivorous invert. | whole body | 2019 | 1 | -28.28 | 4.84 | 49.71 | 10.85 | 4.58 |
| Animal | Invertebrate | *Heteroptera* | Herbivorous invert. | whole body | 2019 | 2 | -27.56 | 5.85 | 50.26 | 10.27 | 4.89 |
| Animal | Invertebrate | *Lumbricidae* | Herbivorous invert. | whole body | 2019 | 2 | -26.85 | 5.28 | 43.11 | 8.91 | 4.84 |
| Animal | Invertebrate | *Symphyta* larvae | Herbivorous invert. | whole body | 2019 | 2 | -29.10 | 3.06 | 48.26 | 7.53 | 6.41 |
| Animal | Invertebrate | *Tettigoniidae* | Herbivorous invert. | whole body | 2019 | 1 | -28.00 | 3.82 | 48.34 | 11.32 | 4.27 |
| Animal | Invertebrate | *Tettigoniidae* | Herbivorous invert. | whole body | 2019 | 2 | -27.75 | 4.39 | 50.49 | 11.03 | 4.58 |
| Animal | Invertebrate | *Araneae* | Predatory invert. | whole body | 2019 | 1 | -26.92 | 7.67 | 45.62 | 11.01 | 4.14 |
| Animal | Invertebrate | *Araneae* | Predatory invert. | whole body | 2019 | 2 | -26.92 | 8.63 | 49.58 | 10.90 | 4.55 |
| Animal | Invertebrate | *Coleoptera* | Predatory invert. | whole body | 2019 | 2 | -26.86 | 7.11 | 50.78 | 10.05 | 5.05 |
| Animal | Invertebrate | *Formicidae* | Predatory invert. | whole body | 2019 | 2 | -27.16 | 6.14 | 50.26 | 10.45 | 4.81 |
| Animal | Invertebrate | *Heteroptera* | Predatory invert. | whole body | 2019 | 1 | -26.76 | 6.24 | 49.09 | 11.32 | 4.34 |
| Animal | Invertebrate | *Heteroptera* | Predatory invert. | whole body | 2019 | 2 | -27.17 | 5.89 | 47.98 | 11.48 | 4.18 |
| Animal | Mammal | *Myodes* | Mammal | brain, muscle | 2019 | | -25.22 | 4.71 | 41.32 | 10.63 | 3.89 |
| Animal | Mammal | *Soricidae* | Mammal | muscle | 2019 | | -24.98 | 6.28 | 41.12 | 11.23 | 3.66 |
| Fungi | Fungi | *Fungi* | Fungi | fruiting body | 2020 | 2 | -26.63 | 5.19 | 45.13 | 4.58 | 9.86 |
| Fungi | Fungi | *Fungi* | Fungi | fruiting body | 2020 | 2 | -25.22 | 7.37 | 41.26 | 6.66 | 6.20 |
| Fungi | Fungi | *Fungi* | Fungi | fruiting body | 2020 | 2 | -24.78 | 3.80 | 45.21 | 4.80 | 9.43 |
| Plant | Forb | *Aegopodium* | Reproductive | inflorescence | 2020 | 1 | -28.47 | 1.37 | 42.11 | 3.89 | 10.83 |
| Plant | Forb | *Aegopodium* | Reproductive | seed | 2020 | 2 | -28.46 | -0.30 | 51.83 | 3.58 | 14.49 |
| Plant | Forb | *Alchemilla* | Reproductive | inflorescence | 2020 | 2 | -28.77 | 1.26 | 43.42 | 2.62 | 16.57 |
| Plant | Forb | *Anthriscus* | Reproductive | inflorescence | 2019 | 1 | -26.97 | -0.97 | 38.46 | 1.83 | 20.99 |
| Plant | Forb | *Anthriscus* | Reproductive | seed | 2019 | 2 | -27.95 | -2.15 | 50.37 | 3.70 | 13.61 |
| Plant | Forb | *Filipendula* | Reproductive | inflorescence, seed | 2019 | 2 | -28.58 | -0.44 | 45.13 | 1.46 | 30.92 |
| Plant | Forb | *Galeopsis* | Reproductive | inflorescence | 2019 | 1 | -27.19 | 3.46 | 39.70 | 2.03 | 19.54 |
| Plant | Forb | *Galeopsis* | Reproductive | seed | 2019 | 2 | -30.48 | 2.70 | 57.92 | 3.32 | 17.45 |
| Plant | Forb | *Onagraceae* | Reproductive | inflorescence | 2020 | 2 | -28.14 | 2.62 | 45.30 | 1.76 | 25.70 |
| Plant | Forb | *Reproductive* | Reproductive | seed | 2020 | 2 | -28.62 | -0.13 | 41.37 | 5.63 | 7.35 |
| Plant | Forb | *Trifolium* | Reproductive | inflorescence | 2020 | 1 | -29.73 | -0.84 | 45.12 | 2.17 | 20.77 |
| Plant | Forb | *Vicia* | Reproductive | inflorescence | 2020 | 2 | -29.60 | 0.47 | 43.57 | 3.03 | 14.38 |
| Plant | Forb | *Aegopodium* | Vegetative | root, leaf | 2020 | 1 | -28.55 | 0.71 | 39.08 | 2.06 | 18.96 |
| Plant | Forb | *Aegopodium* | Vegetative | root, leaf, inflorescence, seed | 2020 | 2 | -28.69 | 0.33 | 34.87 | 1.12 | 31.09 |
| Plant | Forb | *Alchemilla* | Vegetative | root, leaf | 2020 | 2 | -29.94 | 0.29 | 40.38 | 1.79 | 22.51 |
| Plant | Forb | *Anthriscus* | Vegetative | root | 2019 | 1 | -28.56 | -0.57 | 39.11 | 0.55 | 70.98 |
| Plant | Forb | *Anthriscus* | Vegetative | leaf | 2019 | 1 | -30.08 | -0.71 | 42.76 | 2.24 | 19.13 |
| Plant | Forb | *Anthriscus* | Vegetative | leaf | 2019 | 2 | -30.03 | -1.20 | 40.86 | 1.68 | 24.28 |
| Plant | Forb | *Cirsium* | Vegetative | root, leaf | 2019 | 2 | -25.09 | 1.38 | 43.49 | 2.81 | 15.46 |
| Plant | Forb | *Filipendula* | Vegetative | root, leaf | 2019 | 2 | -29.89 | -0.88 | 45.62 | 1.03 | 44.34 |
| Plant | Forb | *Galeopsis* | Vegetative | root | 2019 | 1 | -28.22 | 0.67 | 37.40 | 1.12 | 33.50 |
| Plant | Forb | *Galeopsis* | Vegetative | leaf | 2019 | 1 | -29.75 | 4.16 | 42.48 | 2.97 | 14.30 |
| Plant | Forb | *Galeopsis* | Vegetative | root | 2019 | 2 | -27.54 | -0.10 | 37.45 | 0.67 | 55.57 |
| Plant | Forb | *Galeopsis* | Vegetative | leaf | 2019 | 2 | -29.39 | 2.99 | 43.88 | 1.57 | 27.91 |
| Plant | Forb | *Onagraceae* | Vegetative | root | 2019 | 1 | -27.69 | 1.43 | 42.00 | 2.45 | 17.15 |
| Plant | Forb | *Onagraceae* | Vegetative | leaf | 2019 | 1 | -27.68 | 1.81 | 45.20 | 3.53 | 12.79 |
| Plant | Forb | *Onagraceae* | Vegetative | root | 2019 | 2 | -28.02 | 1.13 | 42.04 | 1.72 | 24.51 |
| Plant | Forb | *Onagraceae* | Vegetative | leaf | 2019 | 2 | -27.87 | 2.22 | 46.64 | 3.09 | 15.12 |
| Plant | Forb | *Rubus* | Vegetative | root, leaf | 2020 | 2 | -30.68 | -1.29 | 42.79 | 1.99 | 21.46 |
| Plant | Forb | *Taraxacum* | Vegetative | root, leaf | 2019 | 2 | -30.44 | 0.77 | 41.51 | 2.36 | 17.58 |
| Plant | Forb | *Taraxacum* | Vegetative | root | 2020 | 2 | -29.56 | -0.51 | 35.70 | 1.12 | 31.91 |
| Plant | Forb | *Taraxacum* | Vegetative | leaf | 2020 | 2 | -30.48 | 0.98 | 40.93 | 3.45 | 11.88 |
| Plant | Forb | *Trifolium* | Vegetative | root, leaf, inflorescence | 2019 | 2 | -29.08 | -0.90 | 44.98 | 3.25 | 13.83 |
| Plant | Forb | *Trifolium* | Vegetative | root, leaf, inflorescence | 2019 | 2 | -28.92 | -1.79 | 44.24 | 2.43 | 18.17 |

| Kingdom | Group | Taxon | Type | Parts | Year | Round | d13C | d15N | %C | %N | C/N |
|---|---|---|---|---|---|---|---|---|---|---|---|
| Plant | Forb | *Tussilago* | Vegetative | root, leaf | 2020 | 1 | -29.92 | 1.26 | 39.73 | 2.19 | 18.15 |
| Plant | Forb | *Urtica* | Vegetative | root, leaf, inflorescence | 2020 | 1 | -27.02 | 1.85 | 40.24 | 3.27 | 12.32 |
| Plant | Forb | *Urtica* | Vegetative | root, leaf, inflorescence, seed | 2020 | 2 | -27.90 | 1.97 | 39.81 | 1.89 | 21.07 |
| Plant | Forb | *Vicia* | Vegetative | root, leaf | 2020 | 2 | -29.61 | 0.15 | 41.58 | 3.05 | 13.61 |
| Plant | Grass | *Agrostis* | Reproductive | inflorescence | 2019 | 1 | -27.97 | 0.04 | 40.10 | 1.47 | 27.33 |
| Plant | Grass | *Elytrigia* | Reproductive | inflorescence | 2019 | 1 | -25.82 | 1.12 | 42.45 | 1.59 | 26.67 |
| Plant | Grass | *Elytrigia* | Reproductive | inflorescence | 2019 | 2 | -26.18 | 1.59 | 42.82 | 0.97 | 44.26 |
| Plant | Grass | *Elytrigia* | Reproductive | seed | 2020 | 2 | -26.83 | 1.31 | 44.42 | 1.03 | 43.10 |
| Plant | Grass | *Phleum* | Reproductive | inflorescence | 2019 | 1 | -26.02 | 2.19 | 45.44 | 1.59 | 28.55 |
| Plant | Grass | *Phleum* | Reproductive | seed | 2019 | 2 | -25.76 | 0.15 | 42.58 | 2.37 | 17.96 |
| Plant | Grass | *Phleum* | Reproductive | inflorescence, seed | 2020 | 2 | -26.74 | 0.28 | 40.80 | 1.30 | 31.35 |
| Plant | Grass | *Poa* | Reproductive | inflorescence | 2019 | 1 | -27.54 | 2.98 | 42.85 | 1.10 | 38.89 |
| Plant | Grass | *Poa* | Reproductive | inflorescence | 2019 | 2 | -28.23 | -0.54 | 40.20 | 0.78 | 51.45 |
| Plant | Grass | *Poaceae* | Reproductive | inflorescence, seed | 2019 | 2 | -25.75 | 0.00 | 41.55 | 1.96 | 21.25 |
| Plant | Grass | *Agrostis* | Vegetative | leaf | 2019 | 1 | -29.58 | -0.18 | 41.48 | 1.74 | 23.79 |
| Plant | Grass | *Dactylis* | Vegetative | root, leaf | 2020 | 2 | -28.33 | 0.60 | 41.42 | 1.33 | 31.09 |
| Plant | Grass | *Elytrigia* | Vegetative | root | 2019 | 1 | -27.83 | 0.53 | 40.46 | 1.03 | 39.38 |
| Plant | Grass | *Elytrigia* | Vegetative | leaf | 2019 | 1 | -28.69 | 2.02 | 38.81 | 1.38 | 28.03 |
| Plant | Grass | *Elytrigia* | Vegetative | root | 2019 | 2 | -27.43 | 1.91 | 38.54 | 0.93 | 41.51 |
| Plant | Grass | *Elytrigia* | Vegetative | leaf | 2019 | 2 | -28.18 | 1.54 | 43.07 | 1.42 | 30.36 |
| Plant | Grass | *Elytrigia* | Vegetative | root, leaf | 2020 | 2 | -31.71 | 0.67 | 40.80 | 2.46 | 16.58 |
| Plant | Grass | *Phleum* | Vegetative | root | 2019 | 1 | -27.84 | -0.50 | 39.92 | 1.06 | 37.62 |
| Plant | Grass | *Phleum* | Vegetative | leaf | 2019 | 1 | -29.06 | 1.10 | 43.32 | 1.44 | 30.03 |
| Plant | Grass | *Phleum* | Vegetative | root, leaf | 2020 | 1 | -28.76 | 2.56 | 33.59 | 1.23 | 27.32 |
| Plant | Grass | *Phleum* | Vegetative | root | 2019 | 2 | -27.47 | -0.06 | 41.87 | 0.72 | 58.21 |
| Plant | Grass | *Phleum* | Vegetative | leaf | 2019 | 2 | -28.74 | -0.26 | 44.23 | 0.80 | 55.37 |
| Plant | Grass | *Poa* | Vegetative | leaf | 2019 | 1 | -29.35 | 4.38 | 42.36 | 0.71 | 59.46 |
| Plant | Grass | *Poa* | Vegetative | root | 2019 | 2 | -29.12 | -0.46 | 40.04 | 0.90 | 44.51 |
| Plant | Grass | *Poaceae* | Vegetative | root, leaf | 2019 | 2 | -29.33 | -0.31 | 40.08 | 0.68 | 59.26 |
| Plant | Moss | *Bryobionta* | Vegetative | whole plant | 2020 | 2 | -32.90 | -2.82 | 43.59 | 1.46 | 29.93 |
| Plant | Moss | *Bryobionta* | Vegetative | whole plant | 2020 | 2 | -29.68 | 0.63 | 44.36 | 1.34 | 32.99 |
| Plant | Moss | *Bryobionta* | Vegetative | whole plant | 2020 | 2 | -27.90 | 1.99 | 29.91 | 1.04 | 28.77 |
| Plant | Lab food | | Rodent chow | pellet | | | -24.908 | 1.89994 | 42.34 | 2.714 | 15.601 |
| Plant | Lab food | | Rodent chow | pellet | | | -25.522 | 1.31336 | 43.3 | 2.906 | 14.899 |
| Plant | Lab food | | Rodent chow | pellet | | | -26.011 | 2.08344 | 43.11 | 2.826 | 15.254 |
| Plant | Lab food | | Rodent chow | pellet | | | -26.016 | 1.94117 | 43.08 | 3.197 | 13.475 |
| Plant | Lab food | | Rodent chow | pellet | | | -25.593 | 1.72642 | 42.89 | 3.043 | 14.097 |
| Plant | Lab food | | Rodent chow | pellet | | | -31.63 | 1.70454 | 37.88 | 1.617 | 23.429 |

*Legend:*

| **Column** | **Content** |
|---|---|
| Kingdom | Grouping of food sources at kingdom level (animal, fungi, plant) |
| Group | Sub-grouping food sources |
| Taxon | Taxonomic division |
| Type | Grouping of food sources according to predicted trophic level |
| Parts | Parts of organism sampled |
| Year | Year of collection (2019 from Peltokangas enclosures, 2020 from Pukara enclosures) |
| Round | Time of sample collection: 1= early summer (June/July), 2= late summer (August/September) |
| d13C | $\delta^{13}$C isotope ratio |
| d15N | $\delta^{15}$N isotope ratio |
| %C | % of carbon in the sample |
| %N | % of nitrogen in the sample |
| C/N | Ratio of nitrogen to carbon in the sample (%C/%N)) |
