## Supplementary information 1 for "Artificial selection for predatory behavior results in dietary niche differentiation in an omnivorous mammal"

### Supplementary methods

#### Food source sampling

To ensure that the isotope ratios in vole hair reflect the isotope ratios in the sources of food available in the field enclosures, we analysed isotopes of carbon and nitrogen also in numerous possible foods collected from the field enclosures, and in the lab pellets routinely fed to voles in captivity. Details on the food sources are provided in Supplementary information 2.

In the summer of 2019 and 2020, we collected samples of plants, fungi and animals that may be consumed by the voles in the field enclosures. Thus, the food source sampling was done in different years than the vole hair sampling, but the timing of the food source sampling matched the season of the early and late season rounds of the experiment so that the sampled items would be representative of the diet available to the voles in each round. We have no reason to assume that there were substantial differences in the isotope composition of the common species in the enclosures between these years. Plant and animal samples were collected in 2019 from the same field enclosures. In 2020 we collected plants and fungi from another nearby set of enclosures for the purposes of another experiment and used a subset of those samples to supplement the range of species found in the 2019 collection. Samples of rodent pellets available to voles in laboratory conditions were collected from the captive colony on six occasions in 2019-2021.

Plant sampling was done by selecting the most common plants (typically covering >10% of the plot) from a 1 $m^2$ square at a random location in each of the 11 enclosures used in the experiment. Entire plants were collected and stored at -20 °C. Fungi were collected from the sampling plots opportunistically. Invertebrate sampling was done in early and late summer from the vegetation with a butterfly net passed through the vegetation along the diagonal of each enclosure. We additionally collected invertebrates by placing one pitfall trap (plastic cup with water and a drop of detergent sunk into the ground to the brim of the cup) in each enclosure for a minimum of 8 h. All samples were stored in plastic containers at -20 °C until sorting and identification and analyses.

Samples were pooled to represent the relevant dietary groups. Plants were identified at least to the family level and where available, vegetative and reproductive parts were separated for the isotope runs. Several plant individuals from the same site and round were pooled into the same samples for the isotope runs. Where the same species of plant was sampled on multiple occasions (different seasons, years, sites), an average was computed of the isotope values of each species, and these were used to compute the mean value for the group to reduce the bias created by sampling some species multiple times and others only once. Invertebrates were identified with a microscope to at least family level (or as needed to determine trophic level: herbivorous vs. omnivorous/carnivorous). Invertebrates belonging to the same taxonomic group from different enclosures were combined into one sample. Identification of fungi was not attempted, instead a representative set of fruiting bodies of different fungi morphotypes was sampled and samples from the same enclosure were pooled.

All samples were freeze dried. Desiccated samples were powdered and 0.95-1.05 mg of each plant and fungi sample and 0.55-0.65 mg of each animal sample was weighed and transferred to tin capsules for stable isotope analysis. Birch (*Betula pubescens*) leaf was used as a standard for the plant samples and pellets and pike (*Esox lucius L.*) white muscle was used for the invertebrate samples.

##### Exploratory statistical analyses

We tested the associations of the isotope ratios and morphological traits with selection regime, season with Wilcoxon tests, and covariates with Kruskal-Wallis and Dunn post-hoc tests and simple linear models, see supplementary results below.

#### Supplementary results

##### Morphological traits

Predatory line offspring tended to be structurally larger (mean head width of Control voles=12.53 mm, Predatory voles=12.76 mm; t = -2.606, df = 128.95, P= 0.010) and slightly heavier than Control-line voles (mean body mass of Control voles= 13.05 g, Predatory voles=13.87 g, t = -1.962, df = 129.33, P= 0.052) but the lines did not differ in body condition (t = 0.532, df = 128.07, P= 0.595). Multiple regression models confirmed an effect of selection regime on structural size (β= 0.293, SE= 0.093, t= 3.164, P= 0.002) and body mass (β=1.339, SE= 0.436, t= 3.071, P= 0.003) but no significant effect of $\delta^{15}$N, $\delta^{13}$C, sex, or replicate round (all P>0.1). Residual body condition was not significantly associated with $\delta^{15}$N, $\delta^{13}$C, sex, replicate round, nor selection regime (linear model of body condition, all predictor variables P>0.1).

##### Average isotope ratios

The raw data indicated that the $\delta^{13}$C values were higher in the late summer (Wilcoxon test W = 260, P<0.001) while $\delta^{15}$N was higher in the early season (W = 3119, P<0.001). Predatory voles had significantly higher $\delta^{13}$C and $\delta^{15}$N-values than control voles in the early season ($\delta^{13}$C: W = 47, P<0.001; $\delta^{15}$N: W = 79, = 0.002), but no such differences were observed in the late season ($\delta^{13}$C: W = 1010, P= 0.529; $\delta^{15}$N: W = 894, P= 0.732; see main text Figure 1).

##### Random effects

Mother's identity (random effect, N=52) had a significant influence on both $\delta^{13}$C ($X^2$=111.02, df = 51, P< 0.001) and $\delta^{15}$N ($X^2$=108.89, df = 51 P< 0.001). There were average differences also among enclosures in $\delta^{15}$N ($X^2$=33.894, df = 9, P< 0.001). The differences in marginal and conditional pseudo-$R^2$ (Table S1) suggest that the random effects of enclosure and mother's id together explained ca. 48% of the variation in $\delta^{15}$N. Mother's id explained ca. 25 % of the variation in $\delta^{13}$C. The average number of juveniles per mother did not significantly differ between tactics (on average 2.6 juveniles with the same mother in C, 2.5 in P).

#### Supplementary tables

Table S1. Estimates from mixed-effects models of the isotope ratios a) $\delta^{15}N$ and b) $\delta^{13}C$ in the hair of juvenile voles at the end of the field experiment. N=132 for both models. “Ref.” indicates the reference level for categorical variables. Significant effects are in bold.

| | **Predictor variable** | **Estimate** | **SE** | **t** | **P** |
|---|---|---|---|---|---|
| a) $\delta^{15}N$ (Box-Cox λ=-1) | Intercept | 0.834 | 0.004 | 192.476 | <0.001 |
| | Round, 2 (ref. 1) | -0.006 | 0.004 | -1.555 | 0.126 |
| | **Selection regime, P (ref. C)** | **0.012** | **0.004** | **2.760** | **0.008** |
| | **Round x Selection** | **-0.013** | **0.005** | **-2.473** | **0.017** |
| | **Density, low (ref. high)** | **0.008** | **0.003** | **2.371** | **0.023** |
| | Sex, male (ref. female) | <0.001 | 0.001 | 0.204 | 0.839 |
| | Body condition | <0.001 | 0.001 | -0.494 | 0.622 |
| b) $\delta^{13}C$ (Box-Cox λ=2) | Intercept | 1.275 | 0.398 | 3.204 | 0.002 |
| | **Round, 2 (ref. 1)** | **3.779** | **0.427** | **8.843** | **<0.001** |
| | **Selection regime, P (ref. C)** | **1.756** | **0.465** | **3.776** | **<0.001** |
| | **Round x Selection** | **-1.987** | **0.569** | **-3.494** | **0.001** |
| | Density, low (ref. high) | 0.176 | 0.270 | 0.652 | 0.518 |
| | Sex, male (ref. female) | -0.143 | 0.152 | -0.940 | 0.349 |
| | Body condition | -0.070 | 0.075 | -0.922 | 0.358 |

a) Random effect: Dam ID (N=52): SD=0.006, Enclosure (N=10): SD=0.007; Residual: SD=0.006. $R^2$marginal=0.348, $R^2$conditional=0.823.

b) Random effect: Dam ID (N=52): SD=0.805, Residual: SD=0.729. $R^2$marginal=0.566, $R^2$conditional=0.804.

Table S2. Predictors of a) $\delta^{15}N$ and b) $\delta^{13}C$ values in predatory and control-line bank voles based on a linear mixed model of all data (untransformed and not excluding an outlier, N=133). Results are shown for a model combining both replicates and for the replicates separately. "Ref." indicates the reference level for categorical variables. Statistically significant results in bold.

| | **Predictor variable** | **Estimate** | **SE** | **t** | **P** |
|---|---|---|---|---|---|
| a) $\delta^{15}N$ | Intercept | 6.029 | 0.184 | 32.729 | <0.001 |
| | Round (ref. 1) | -0.223 | 0.179 | -1.249 | 0.217 |
| | **Selection regime (ref. C)** | **0.453** | **0.194** | **2.328** | **0.024** |
| | Round x Selection | -0.409 | 0.234 | -1.749 | 0.087 |
| | Density (ref. high) | 0.254 | 0.137 | 1.850 | 0.074 |
| | Sex (ref. female) | 0.076 | 0.061 | 1.261 | 0.210 |
| | **Body condition** | **-0.083** | **0.029** | **-2.889** | **0.005** |
| b) $\delta^{13}C$ | Intercept | -25.855 | 0.144 | -179.668 | <0.001 |
| | **Round (ref. 1)** | **1.494** | **0.155** | **9.668** | **<0.001** |
| | **Selection regime (ref. C)** | **0.797** | **0.168** | **4.735** | **<0.001** |
| | **Round x Selection** | **-0.862** | **0.206** | **-4.194** | **<0.001** |
| | Density (ref. high) | 0.100 | 0.098 | 1.028 | 0.310 |
| | Sex (ref. female) | -0.023 | 0.053 | -0.434 | 0.665 |
| | Body condition | -0.033 | 0.026 | -1.277 | 0.204 |

a) Random effect: Dam ID (N=52): SD=0.270, Enclosure (N=10): SD=0.233; Residual: SD=0.298. $R^2$marginal=0.322, $R^2$conditional=0.721.

b) Random effect: Dam ID (N=52): SD=0.296, Residual: SD=0.255. $R^2$marginal=0.595, $R^2$conditional=0.828.

#### Supplementary figures

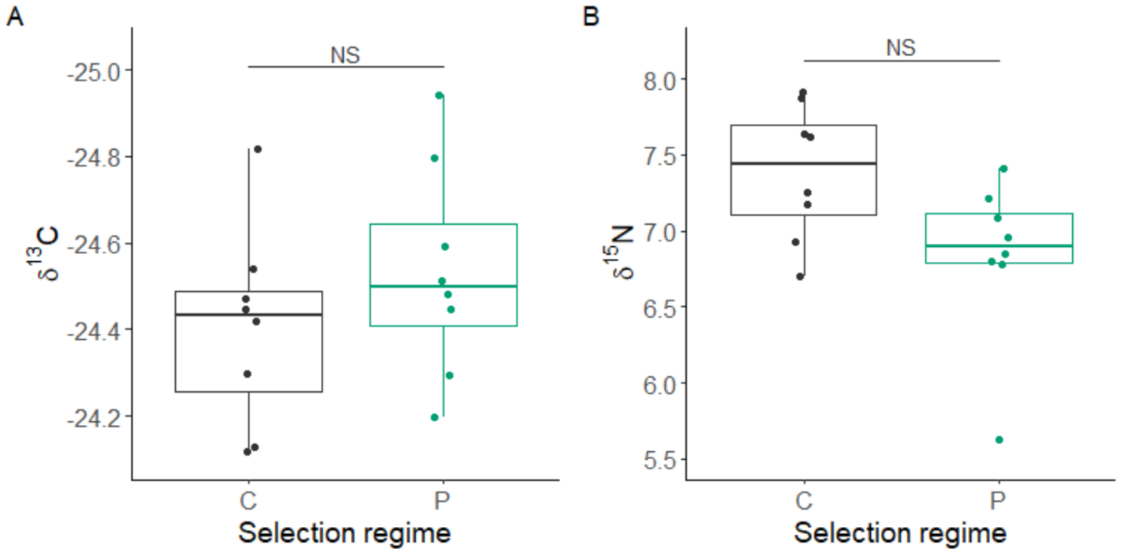


Figure S1. Isotopes of A) $\delta^{13}C$ and B) $\delta^{15}N$ in the hair of captive voles from C- and P-lines (N=16).

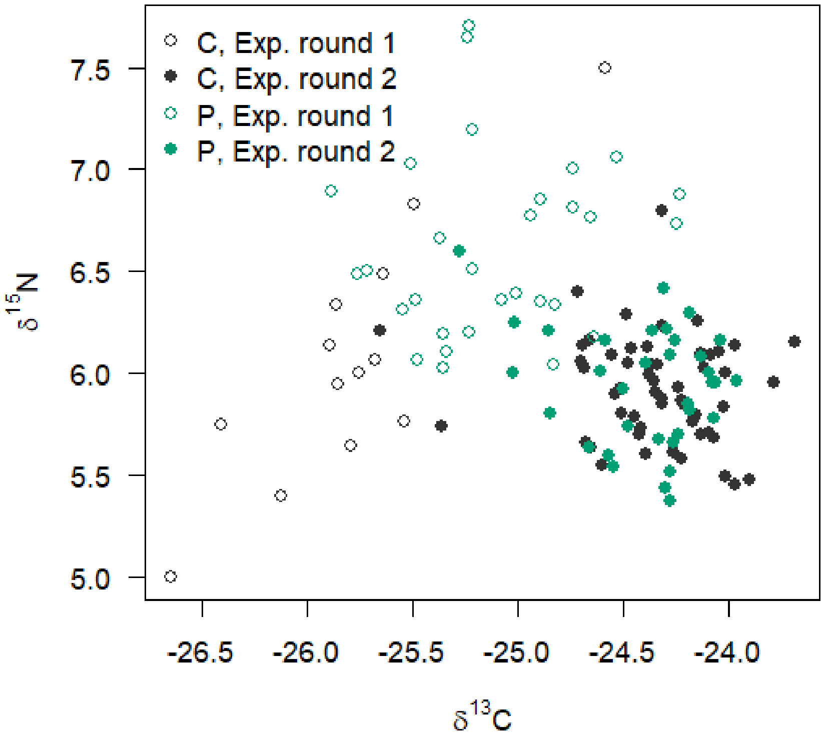


Figure S2. Scatterplot of $\delta^{13}C$ and $\delta^{15}N$ in the hair of C- and P-lines voles in the two experimental rounds.

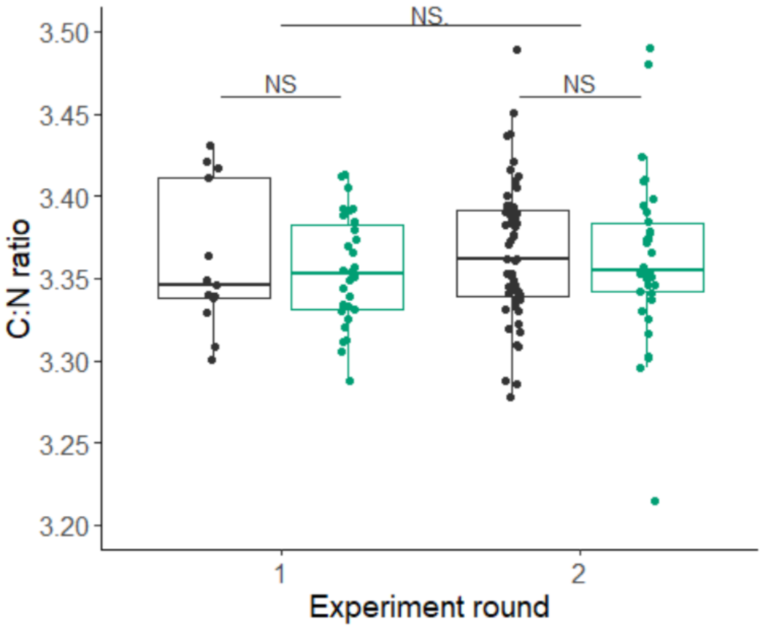


Figure S3: The ratio of carbon to nitrogen in the control (black) and predatory (green) selection regimes and experimental rounds. We found no indication that selection regime and experimental round (or their interaction) would influence the total organic carbon to nitrogen ratio (C:N ratio) in vole hair (after lipids were removed), with the variation among samples being overall very small.
